# The neutral frequency spectrum of linked sites

**DOI:** 10.1101/100123

**Authors:** Luca Ferretti, Alexander Klassmann, Emanuele Raineri, Sebastián E. Ramos-Onsins, Thomas Wiehe, Guillaume Achaz

## Abstract

We introduce the conditional Site Frequency Spectrum (SFS) for a genomic region linked to a focal mutation of known frequency. An exact expression for its expected value is provided for the neutral model without recombination. Its relation with the expected SFS for two sites, 2-SFS, is discussed. These spectra derive from the coalescent approach of Fu (1995) for finite samples, which is reviewed. Remarkably simple expressions are obtained for the linked SFS of a large population, which are also solutions of the multiallelic Kolmogorov equations. These formulae are the immediate extensions of the well known single site *θ*/*f* neutral SFS. Besides the general interest in these spectra, they relate to relevant biological cases, such as structural variants and introgressions. As an application, a recipe to adapt Tajima’s *D* and other SFS-based neutrality tests to a non-recombining region containing a neutral marker is presented.

## 1. Introduction

One of the basic features that characterizes nucleotide polymorphisms is the Site Frequency Spectrum (SFS), that is the distribution of the mutation frequencies at each site. The SFS can be computed either for the whole (large) population, assuming that the frequency *f* is a continuous value in (0, 1) or for a sample of *n* individuals, for which the frequency is a discrete variable *f* = *k*/*n*, where *k ∈* [1, *n* – 1]. Sites with alleles at frequency 0 or 1 are not included in the SFS.

According to the standard neutral model of molecular evolution [1], polymorphisms segregating in a population eventually reach a mutation-drift equilibrium. In this model, the expected neutral spectrum is proportional to the inverse of the frequency [2, 3]. Using coalescent theory, Fu [4] derived the mean and covariance matrix for each component of the sample SFS, by averaging coalescent tree realizations across the whole tree space. For a single realization of the coalescent tree, results are different and depend on the realization; for example, mutations of high frequencies can be present only for highly unbalanced genealogies [5]. The SFS was also studied in scenarios including selection [6, 7], demography [8, 9] or population structure [10].

Besides its general interest, the SFS has been used to devise goodness-of-fit statistical tests to estimate the relevance of the standard neutral model for an observed dataset. SFS-based neutrality tests contrast estimations of the nucleotide variability from different bins of the sample SFS [11, 12, 13]. It was shown that, once the SFS under an alternative scenario (e.g. selection, demography or structure) is known, the optimal test to reject the standard neutral model is based on the difference between the standard neutral SFS and the alternative scenario SFS [14]. All these tests assume complete linkage among variants in their null model.

Assuming independence between the sites, the observed SFS can also be used to estimate model parameters. An interesting recent approach is the estimation of piece-wise constant demography from genomewide SFS (e.g. [15]). More sophisticated methods based on the expected SFS, such as Poisson Random Field [16, 17, 18] and Composite Likelihood approaches [*e.g*., 7, 19, 20, 21], have also played an important role in the detection of events of selection across regions of the genome. However, the assumption of linkage equilibrium is often violated in genetic data. In fact, while the average spectrum is insensitive to recombination, the knowledge on linked variants affects the distribution of summary statistics, therefore the spread (and possibly the mean) of the estimated parameters [22, 23]. For this reason, simulations of the evolution of linked sequences are required for an accurate estimation of the statistical support for different models [24].

The joint SFS for multiple sites has been the subject of longstanding investigations. The simplest spectrum for multiple sites is the “two-locus frequency spectrum” [25], which we name the “two-Sites Frequency Spectrum” or 2-SFS. Assuming independence between the sites (*i.e*. free recombination), it simply reduces to the random association between two single-sites spectra (1-SFS). For intermediate recombination, a recursion solvable for small sample size has been provided [26, 27] as well as a numerical solution relying on simulations [25]. Even without recombination, finding an analytical expression for the spectrum has proven to be difficult.

There is a close relation between the *m*-SFS (the joint SFS of *m* sites) and the multi-allelic spectrum of a single *locus* (defined as a sequence with one or more sites). Under the *infinite-sites* model, sites are assumed to have at most two alleles as new mutations occur exclusively at non-polymorphic sites. At the locus scale, each haplotype (the specific combination of the alleles carried at each locus) can be interpreted as a single allele at a multi-allelic locus. In the absence of recombination, each point mutation either leaves the number of different haplotypes unchanged or generates one new haplotype. Therefore, at least conceptually, the SFS for *m* non-recombinant biallelic sites at low mutation rate is closely related to the spectrum of *m* + 1 alleles in a multi-allelic locus. Indeed, it is possible to retrieve the latter from the former by considering the *m* + 1 alleles that result from the *m* polymorphic sites. However, the *m*-SFS contains extra-information on the different couplings between sites that is not available in the multi-allelic spectrum.

For an infinite population, the multi-alleles single-locus spectrum is the solution of a multiallelic diffusion equation [3, section 5.10]. Polynomial expansions were proposed to solve the diffusion equations for the SFS of an infinite population [28, 29, 30], as well as moment-based approaches [31]. Finally, a polynomial expansion of the 2-SFS has been found for two sites without recombination and with general selection coefficients [32]. However, the reported solution is an infinite series and is in sharp contrast with the simplicity of the solution for a single neutral site: E[*ξ*(*f*)] = *θ*/*f*. Furthermore, no closed form was provided for the 2-SFS of a sample.

Using a coalescent framework, the probability and size of two nested mutations were expressed by Hobolth and Wiuf [33] as sums of binomial coefficients. Their formulae can be rewritten as an expected SFS in terms of a finite series. However their conditioning on exactly two nested mutations skews the spectrum and simulations show that their result is valid only for *Lθ* ≪ 1. Interesting analytical results on the spectrum of tri-allelic loci and recurrent mutations were obtained by Jenkins, Song and collaborators [34, 35] for the Kingman coalescent and general allelic transition matrices. More recently, Sargsyan [36] generalized the result of [33] by conditioning on any two mutations (nested or not) and extending it to populations of variable size. Moreover, he clarified the notion and classification of the 2-SFS.

In this work, we review and present in its simplest possible form the exact solution for the expectation of the neutral sample 2-SFS without recombination, then we extend it to a closed-form solution for the continuous population 2-SFS. The solution for a finite sample was derived previously in many disguises in a coalescent framework [4, 34, 37, 36] and its extrapolation to the limit of infinite sample sizes yields the continuous spectrum, which is a solution of the multi-allelic Kolmogorov equations. Furthermore, we derive the expected 1-SFS of sites that are completely linked to a focal mutation of known frequency. This spectrum has several potential applications. In section S1 of the Supplementary Material we extend the formulae for the continuous 2-SFS to closed expressions for the multi-allelic spectrum of a locus with three alleles.

Finally, as an application, we present a recipe to build a class of SFS-based neutrality tests for sequences containing a known neutral marker of given frequency. This is a typical scenario when the marker (and the region around it) has been detected independently as an outlier in genome-wide association studies or population differentiation studies with SNP arrays. As far as we know, this is the first proper adaptation of Tajima’s *D* and similar statistical tests to this kind of sequence data.

### Model definition and notation

We consider a population of *N* haploid individuals without recombination. All subsequent results can be applied to diploids, provided that 2*N* is used instead of *N*, and to other cases by substituting the appropriate effective population size. We denote by *µ* the mutation rate per site and by *θ* = 2*Nµ* the population-scaled mutation rate per site. We work in the infinite-sites approximation, that is valid in the limit of small mutation rates *θ* ≪ 1. More precisely, our results are derived in the limit *θ* → 0 with fixed non-zero *θL*, where *L* is the length of the sequence. The expected value E[.] denotes the expectation with respect to the realizations of the evolutionary process for the sequences in the sample or in the whole population. We use *mutation* as a synonym for derived allele.

### Connection between sample and population SFS

We denote by *ξ*(*f*) the *density* of mutations at frequency *f* in the whole population and by *ξ*_*k*_ the *number* of mutations at frequency *k*/*n* in a sample of size *n*. Importantly, in both cases *f* or *k* refer to the frequency of the mutation, *i.e*. of the *derived* allele, and thus *ξ* corresponds to the *unfolded* SFS.

The two spectra (sample and population) are related. Assuming that a mutation has frequency *f* in the population, the probability of having *k* mutant alleles in a random sample of size *n* is simply given by the Binomial 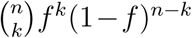. As the expected density of mutations at fixed frequency *f* in the population is given by E[*ξ*(*f*)], one can easily derive the sample frequency from the population frequency using the following sampling formula:

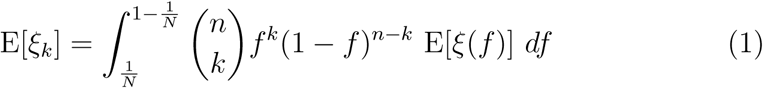

assuming that *n* ≪ *N*.

Conversely, the population SFS can be derived from the sample SFS using the limit of large sample size *n → ∞*. For a sample of *n* individuals, the interval between the frequency bins is 1/*n* and therefore the density of mutations at the continuous frequency *f* = *k*/*n* can be approximated^1^ by 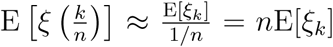. The expected population spectrum can then be constructed from the limit:

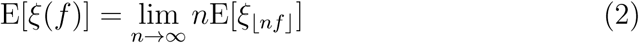

for frequencies not too close to 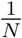 or 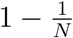.

For a sample of size *n*, the expected neutral spectrum for constant population size is E[*ξ*_*k*_] = *θL*/*k* and consequently, we have E[*ξ*(*f*)] = *θL*/*f* [2, 3]. These results are exact for the Kingman coalescent and the diffusion equations respectively, and they are approximately valid for neutral models for frequencies 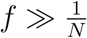. For frequencies of order 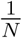, model-dependent corrections are needed and equation (2) is not valid anymore.

In the rest of this section we will deal with sample and population spectra together. We will slightly abuse the notation and switch between number and density of mutations, or probability and probability density.

### Conditional 1-SFS and joint 2-SFS

In the following, we will use two related but different kinds of spectra.

The first kind is the joint 2-SFS of two bi-allelic sites. It is denoted *ξ*(*f*_1_*, f*_2_) for the population and *ξ*_*k,l*_ for the sample. It is defined as the density of pairs of sites with mutation frequencies at *f*_1_ and *f*_2_ for the population (resp. *k*/*n* and *l*/*n* for the sample). This is a natural generalization of the classical SFS for a single site. The expected spectrum E[*ξ*(*f*_1_*, f*_2_)] has multiple equivalent interpretations in the small *θ* limit: (a) for a sequence, it is the expected density of pairs of sites that harbor mutations with frequencies *f*_1_ and *f*_2_; (b) for two randomly chosen linked polymorphic sites, it is the probability density that they contain mutations with frequencies *f*_1_ and *f*_2_. Here we always consider unordered pairs of sites (the ordered case is discussed in section S2).

The second kind of spectrum is a conditional 1-SFS, a frequency spectrum of sites that are linked to a focal mutation of frequency *f*_0_. It is denoted *ξ*(*f*│*f*_0_) for the population and *ξ*_*k*│*l*_ for the sample. Again, this spectrum represents both (a) the expected density of single-site mutations of frequency *f* in a locus linked to a focal neutral mutation of frequency *f*_0_ and (b) the probability density that a randomly chosen site (linked to the focal site) hosts a mutation at frequency *f*.

Note that despite the similarity in notation, the two spectra *ξ*(*f, f*_0_) and *ξ*(*f*│*f*_0_) are different. The difference is the same as the one between the *joint probability p*(*f, f*_0_) that two sites *x* and *x*_0_ have mutations of frequency *f* and *f*_0_ respectively, and the *conditional probability p*(*f*│*f*_0_) that a mutation at site *x* has frequency *f* given that there is a mutation of frequency *f*_0_ at a focal linked site *x*_0_. Furthermore, the joint spectrum *ξ*(*f, f*_0_) refers to pairs of sites – *i.e*. it is a 2-SFS – while the spectrum of linked sites *ξ*(*f*│*f*_0_) is a single-site SFS.

The relation between both types of spectra can be understood from the relation between the probabilities. The expected spectrum E[*ξ*(*f*)] is given by the probability to find a mutation of frequency *f* at a specific site, multiplied by the length of the sequence: E[*ξ*(*f*)] = *p*(*f*)*L*. As noted above, when *L* = 1 (i.e. a locus with a single site is considered), E[*ξ*(*f*)] corresponds to a proper probability *p*(*f*). Assuming the presence of a mutation of frequency *f*_0_ at a focal site, we have E[*ξ*(*f*│*f*_0_)] = *p*(*f*│*f*_0_)(*L* – 1). For pairs of sites, the expected number of mutations at frequencies (*f, f*_0_) is E[*ξ*(*f, f*_0_)] = *p*(*f, f*_0_)*L*(*L* – 1) when *f* ≠ *f*_0_ or *p*(*f*_0_*, f*_0_)*L*(*L* – 1)/2 when *f* = *f*_0_. The additional factor 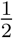 accounts for the symmetrical case of equal frequencies *f* = *f*_0_. The equality *p*(*f, f*_0_) = *p*(*f*│*f*_0_)*p*(*f*_0_) applied to sample and population spectra, results in the following relations:

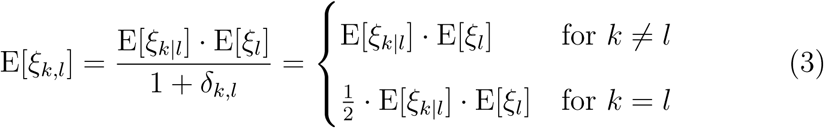

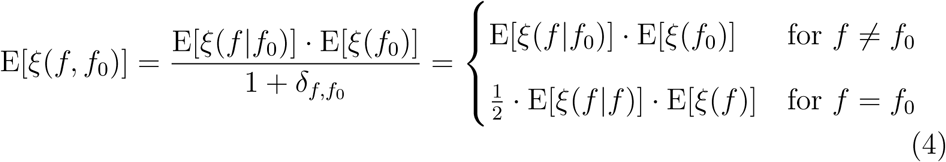

where *δ*_*x,y*_ is 1 if *x* = *y*, and 0 otherwise. Note that *x* and *y* can be either discrete or continuous variables.

By definition, the 2-SFS includes only pairs of sites that are *both* polymorphic. The probability that a pair of sites contains a single polymorphism of frequency *k*/*n* depends only on the 1-SFS and it is approximately equal to 2E[*ξ*_*k*_] for *θ* ≪ 1. Consequently, on a sequence of size *L* hosting *S* polymorphic sites, the number of pairs of sites for which only one of the two is polymorphic of frequency *k*/*n* is E[(*L* – *S*)*ξ*_*k*_] = *L* · E[*ξ*_*k*_] – E[*Sξ*_*k*_] ≈ *L* · E[*ξ*_*k*_] for small *θ*.

## 2. Results

### 2.1 Decomposition of the 2-SFS

We follow [36] and divide the 2-SFS *ξ*(*f*_1_*, f*_2_) without recombination into two different components: one *nested* component *ξ*^(*n*)^(*f*_1_*, f*_2_) for cases where there are individuals carrying the two mutations (one is “nested” in the other), and a *disjoint* component *ξ*^(*d*)^(*f*_1_*, f*_2_) that includes disjoint mutations that are only present in different individuals. The overall spectrum is given by:

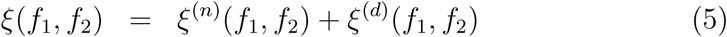

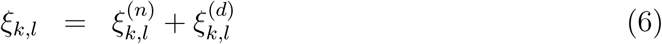

It is noteworthy to mention that the overall spectrum is not sufficient to provide a full description of the genetic state of the two sites, while the two components *ξ*^(*n*)^(*f*_1_*, f*_2_), *ξ*^(*d*)^(*f*_1_*, f*_2_) are enough to reconstruct the genetic content of the two sites up to permutations of all the haplotypes, as it happens with the usual SFS for one site. For example, the following haplotypes (derived alleles marked in bold)

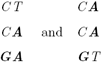

are identical from the point of view of the overall two-loci spectrum: in both samples there is just a pair of mutations with allele count 1 and 2 respectively, therefore the only (symmetrical) nonzero value of the spectrum is *ξ*_1,2_ = *ξ*_2,1_ = 1. However the samples can be distinguished by the two components, since in the first one the mutations are nested 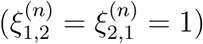, while in the second one they are disjoint 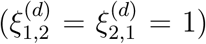. For this reason, these two components constitute the core of the two-loci SFS.

Without recombination, the conditional 1-SFS *ξ*(*f*│*f*_0_) can be also decomposed further^2^ into different subspectra. They are illustrated in Figure 1:

**Figure 1:**
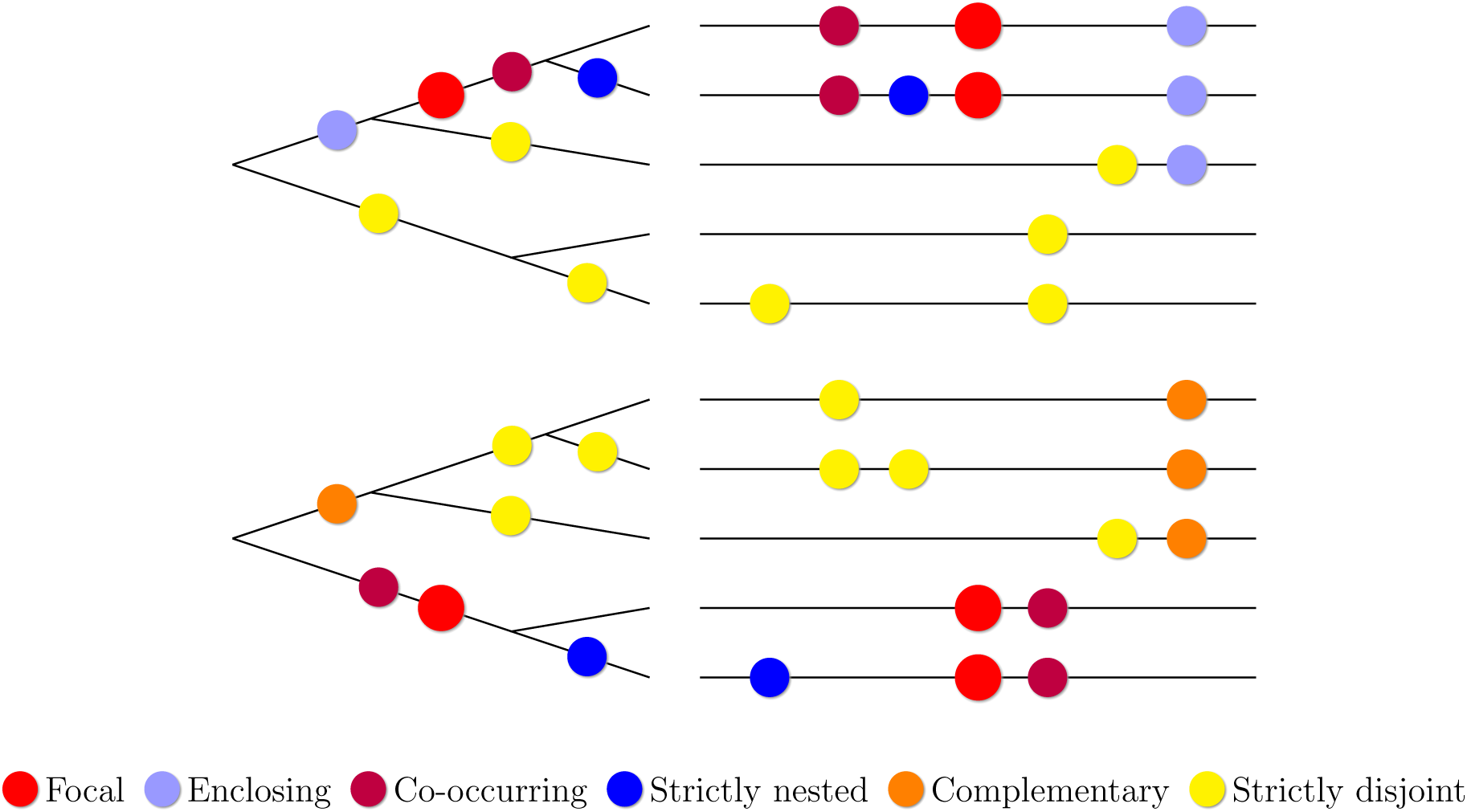
A schema of two non-recombining genomic regions and their corresponding genealogical trees. The black lines on the right represent sequences and the colored circles derived alleles. This figure illustrates the classification of all possible types of mutations with respect to the focal mutation (in red) and their occurrence on the sequence tree. If the focal mutation is not on a root branch (upper panel), it is clear that mutations can be on the same branch as the focal mutation (*co-occurring*), on the subtree below (*strictly nested*), between the focal mutation and the root (*enclosing*), or on other branches (*strictly disjoint*). If the mutation is on a root branch (lower panel), there cannot be enclosing mutations, but there can be mutations on the other root branch (*complementary*).

- *ξ*^(*sn*)^(*f*│*f*_0_): *strictly nested* mutations, where the mutation is carried only by a subset of individuals with the focal mutation;
- *ξ*^(*co*)^(*f*│*f*_0_): *co-occurring* mutations, where both mutations are carried by the same individuals;
- *ξ*^(*en*)^(*f*│*f*_0_): *enclosing* mutations, where only a subset of individuals with the mutation also carry the focal one;
- *ξ*^(*cm*)^(*f*│*f*_0_): *complementary* mutations, where each individual has exactly one of the two mutations;
- *ξ*^(*sd*)^(*f*│*f*_0_): *strictly disjoint* mutations, where the mutation is carried by a subset of the individuals without the focal one.

Importantly, without recombination, enclosing and complementary mutations cannot be present together in the same sequence, as both types of branches are exclusive in a single tree.

With the above definition and using the rules of conditional probabilities *p*(*f, f*_0_) = *p*(*f*│*f*_0_)*p*(*f*_0_) and the interpretations discussed in the previous section, the relations between the two sets of population subspectra are:

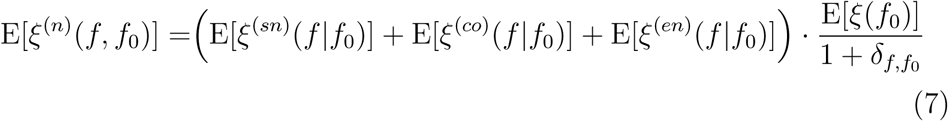

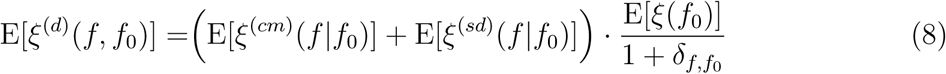

Similarly, for sample spectra, we have

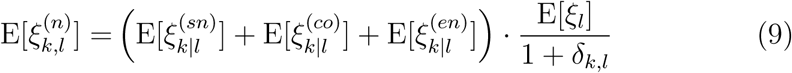

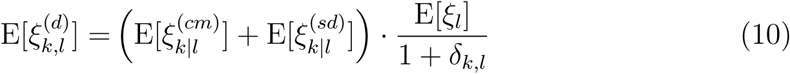

### 2.2 The joint and conditional SFS

In this section, we present the conditional and joint spectra for the sample and the population. The derivations and proofs of all equations in this section are given in Methods and sections S3 and **??** of the Supplementary Material. The folded version of the 2-SFS is provided in Appendix A.

#### 2.2.1 The sample joint 2-SFS

The 2-loci spectrum appeared in the literature under many guises [4, 34, 37, 36]. In the infinite-sites neutral model without recombination, its expected value has a simpler form^3^:

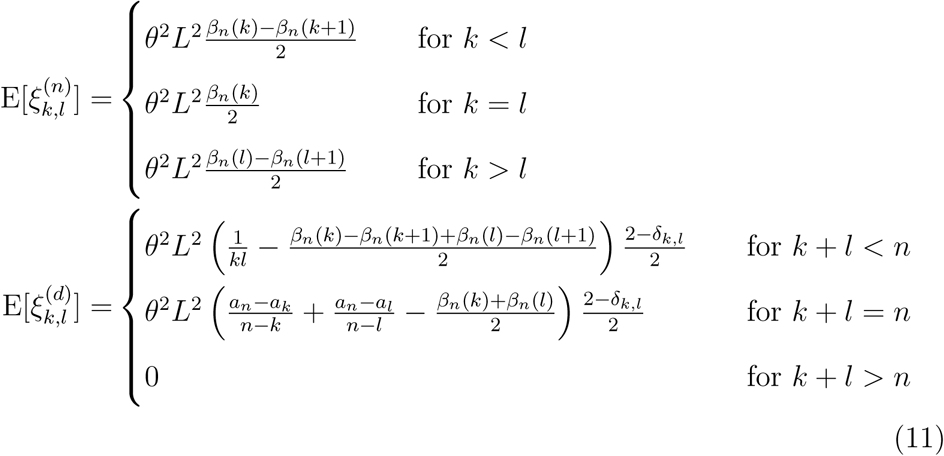

with

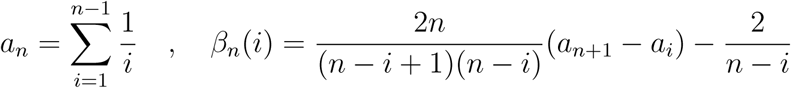

As shown by equation (6), the full spectrum is simply the sum of the two above equations.

#### 2.2.2 The population joint 2-SFS

Similarly, the 2-SFS for the whole population is given by the combination of the two following equations:

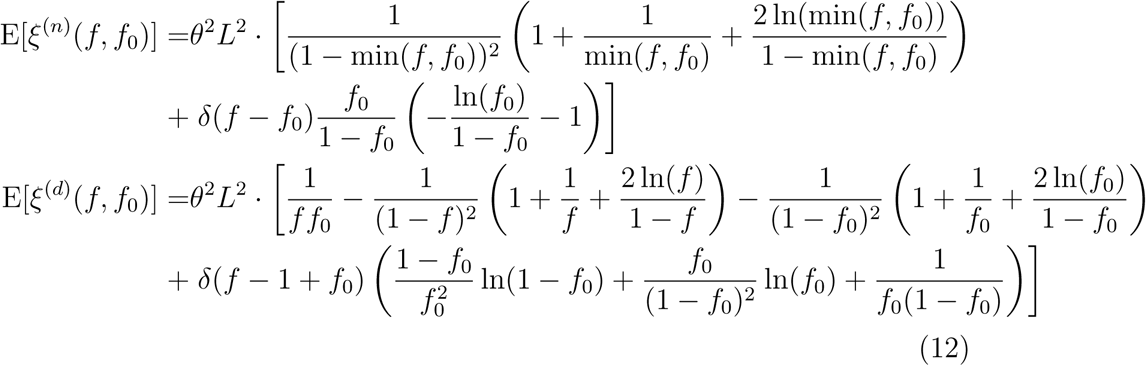

with E[*ξ*^(*n*)^(*f, f*_0_)] = 0 for *f* > *f*_0_ and E[*ξ*^(*d*)^(*f, f*_0_)] = 0 for *f* + *f*_0_ > 1.

Here, we denote by *δ*(*f* – *f*_0_) the density of the Dirac delta distribution concentrated in *f*_0_ (i.e. *δ*(*f* – *f*_0_) = 0 for *f* ≠ *f*_0_, normalized such that 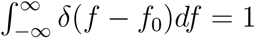.

#### 2.2.3 The sample conditional 1-SFS

The conditional 1-SFS for sites that are linked to a focal mutation of count *l* is simply the sum of all its components, given by the following equations:

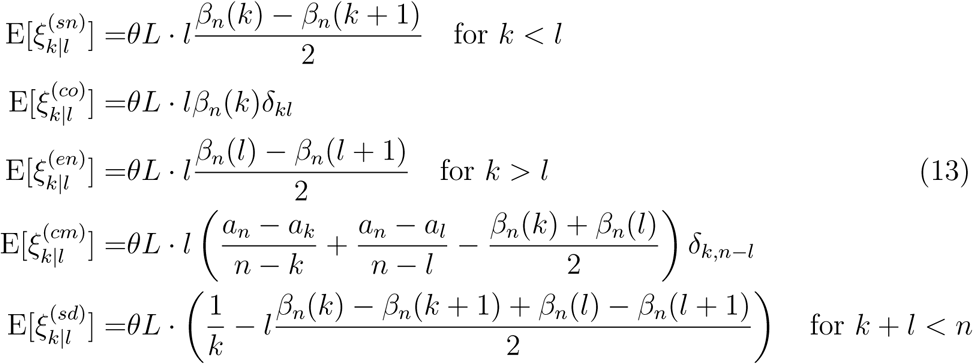

Please note that hereafter unmet conditions imply 0 otherwise.

The strictly nested component of the conditional 1-SFS and its applications have been discussed by Griffiths and Tavare [38].

#### 2.2.4 The population conditional 1-SFS

For the whole population, the expected linked SFS becomes:

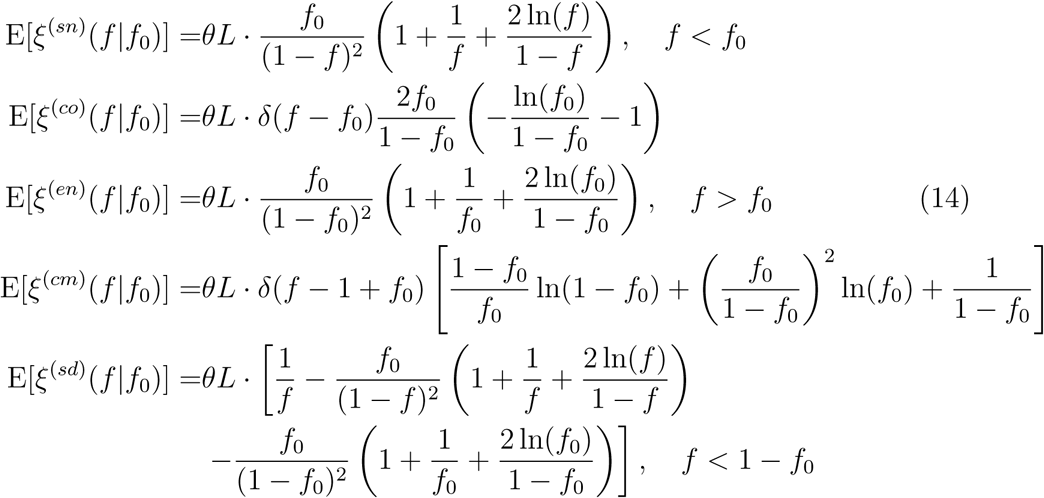

### 2.3 Shape of the SFS

We report the full joint 2-SFS as well as the nested and disjoint components (Figure 2). Nested mutations have preferentially a rare mutation in either site – so that the mutation at lower frequency is easily nested into the other – or are co-occurring mutations. Disjoint mutations are dominated by cases where both mutations are rare, or by complementary mutations. The large contribution of co-occurring (nested component) and complementary mutations (disjoint component) is a direct consequence of the two long branches that coalesce at the root node of a Kingman tree.

**Figure 2:**
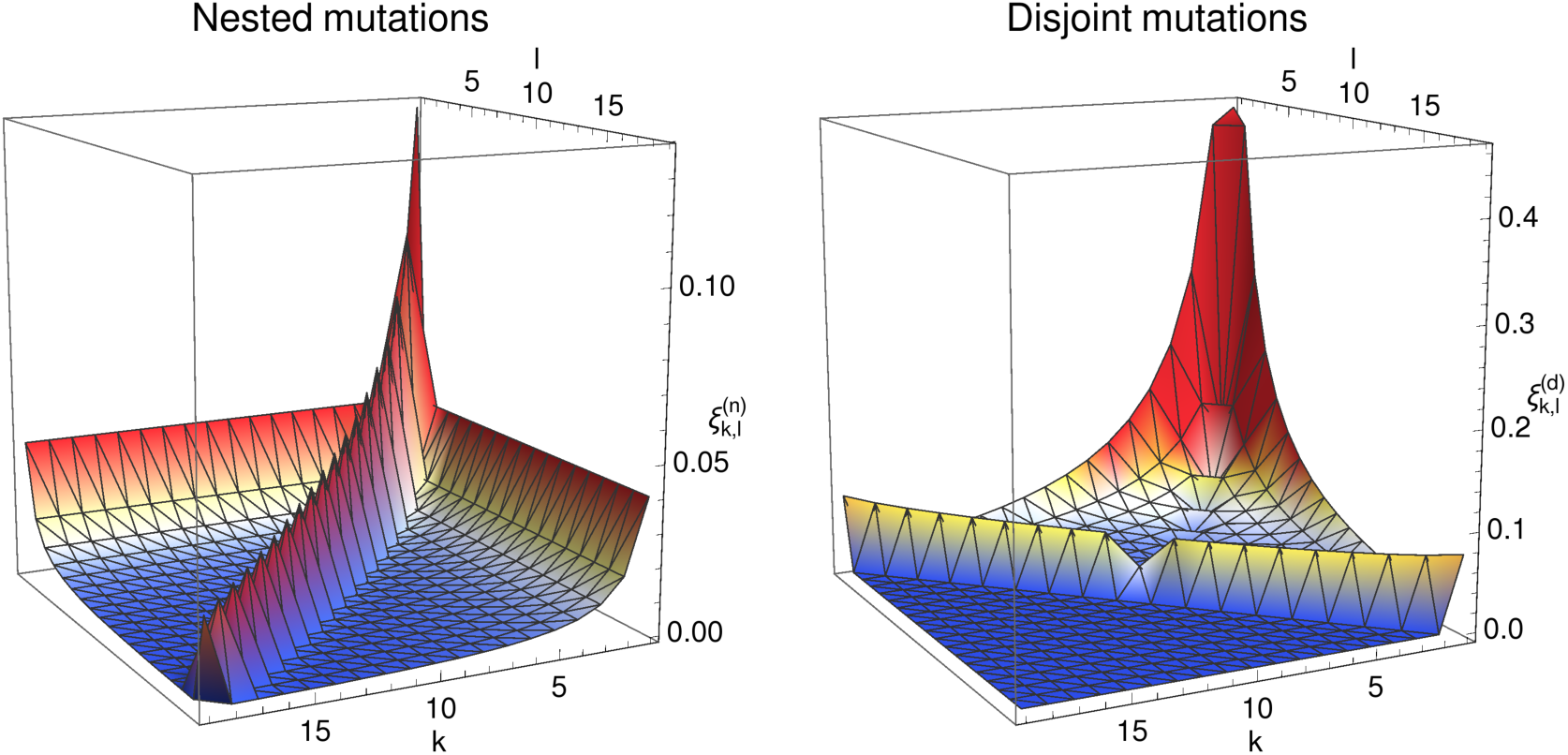
Plots of nested and disjoint contributions to the two-locus frequency spectrum for *θL* = 1, *n* = 20. Note the different scales of the two plots.

The conditional 1-SFS of linked sites and the relative contributions of each component to each frequency are shown in Figure 3. Co-occurring and complementary mutations also account for a considerable fraction of the spectrum, especially when the focal mutation (*f*_0_) is at high frequency. The rest of the spectrum is biased towards mutations with a lower frequency than the focal one. Strictly nested mutations are important only when the frequency of the focal mutation is intermediate or high. Enclosing mutations are rare and their frequencies are uniformly distributed, as previously noted [33].

**Figure 3:**
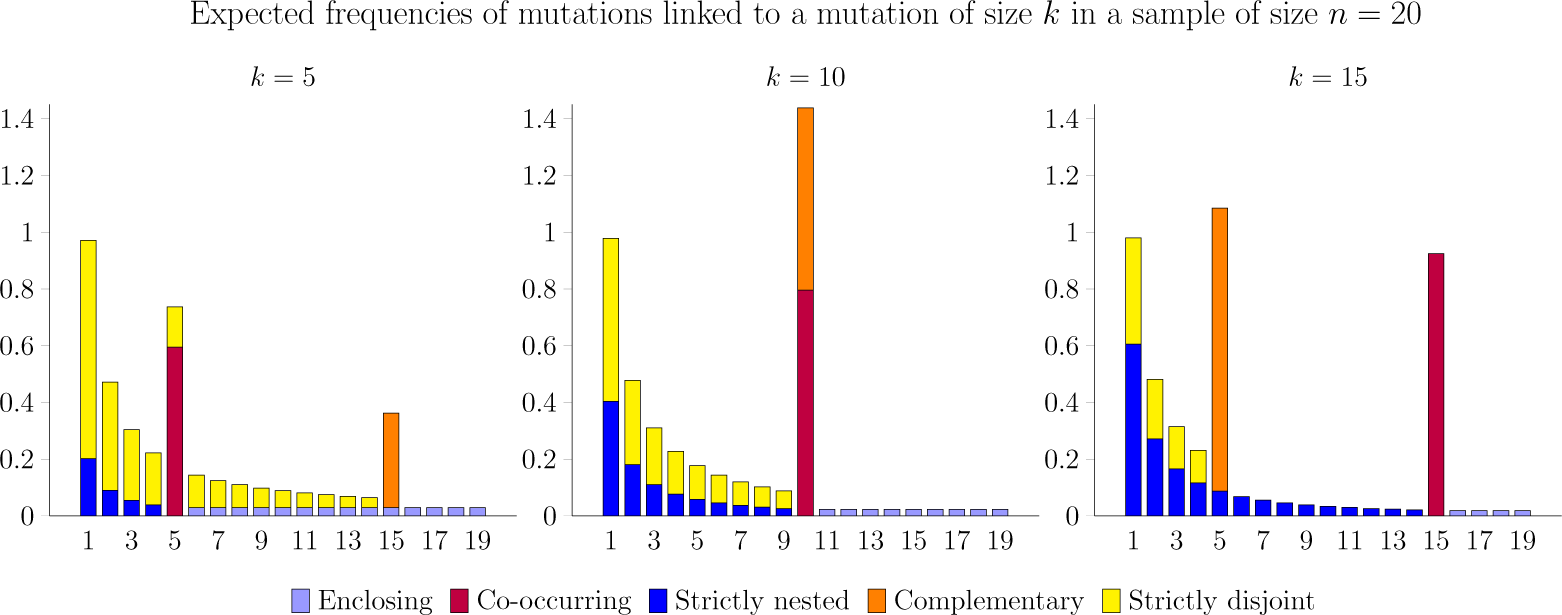
Barplot of the spectrum of linked sites for *θL* = 1, *n* = 20, each column colored according to the different contributions. The focal mutation has frequency 0.25 (left), 0.5 (middle) and 0.75 (right) respectively.

Finally, in Figure 4 we show the impact of a focal mutation of given frequency on two estimators of *θ*. The Watterson’s estimator 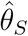 [39] depends on the total number of polymorphic sites, which increases with the frequency of the focal mutation, as they inflate increasingly upper sections of the trees that contribute more to the total tree length. On the other hand, Tajima’s estimator, 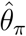 [40] is more sensitive to mutations of intermediate frequency. The difference between the two illustrates how the spectrum is skewed towards common or rare mutations. As Tajima’s *D* [11] is proportional to the difference 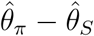, positive values for this test statistic suggest an excess of common mutations while negative values point to an excess of rare mutations. Figure 4 shows that the spectrum has a slight excess of rare mutations at low frequencies of the focal mutation and an excess of common mutations for intermediate frequencies, while it is dominated again by rare mutations if the focal mutation is at high frequencies.

**Figure 4:**
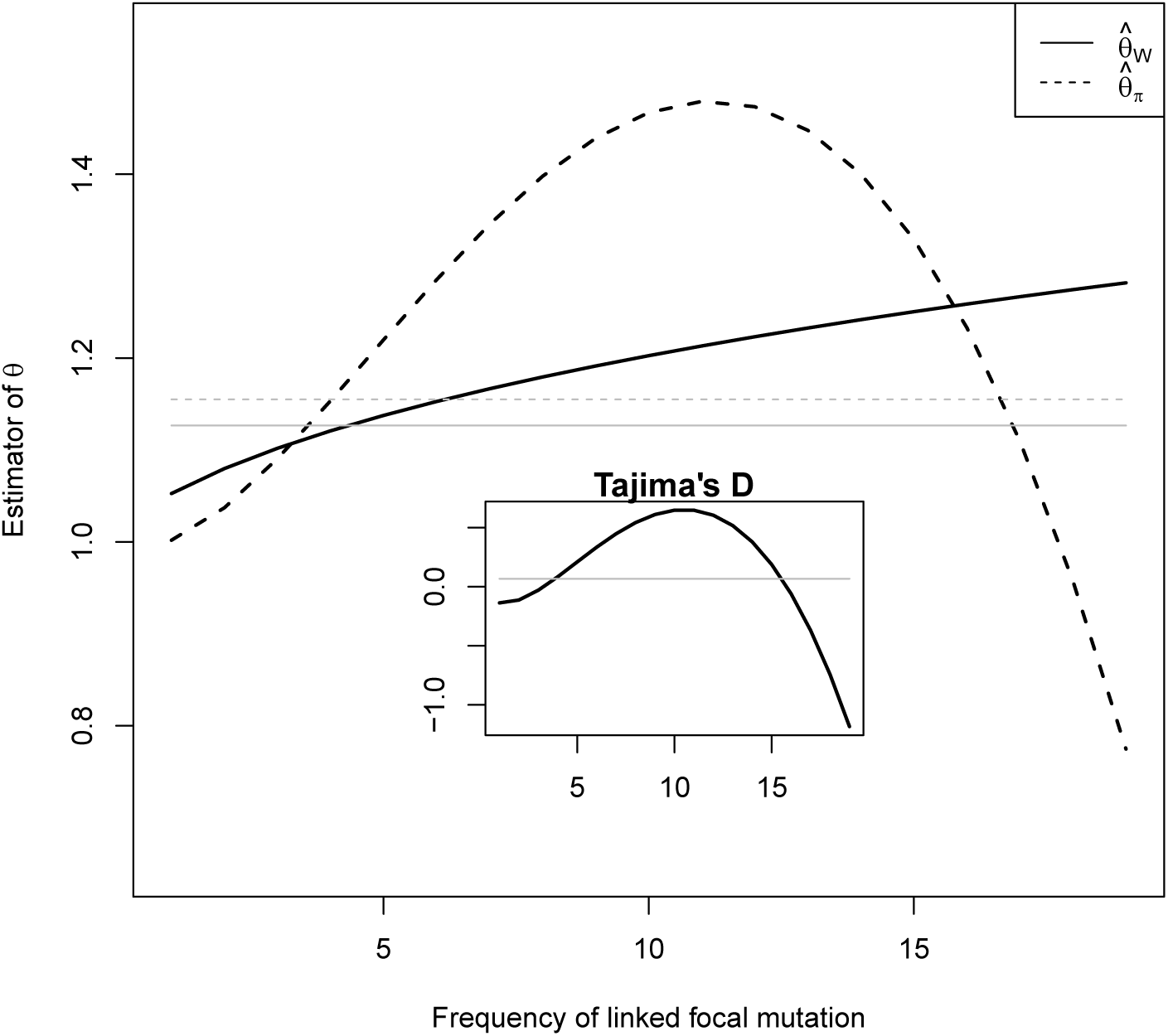
Mean values of the Watterson estimator 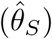 and Tajima estimator 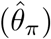 of *θ* conditioned on the presence of a linked mutation, for *θ* = 1, *n* = 20. In the inset, approximate mean value of Tajima’s *D* (computed substituting *S* with its mean value in the denominator). The grey lines represent the expected values conditioned on the presence of a linked polymorphism of any frequency.

### 2.4 Neutrality tests for regions linked to a polymorphic neutral marker

Biallelic putative neutral markers are often used to find regions of interest in a genome. For example, genotype data from SNP arrays can be used together with phenotype measurements to find regions associated with a specific phenotype. Alternatively, if data from multiple populations are available, markers with highly differentiated frequency between populations - i.e. high *F*_*st*_ - can be used to infer potential targets of local selection. It is then natural to use sequence data to test for neutral evolution in a window around the neutral focal marker of known frequency.

Up to now, such tests did not take into account the information given by the frequency of the marker itself. However, since typical markers have biased frequencies towards intermediate values [41], the expected neutral frequency spectrum will likely be dramatically altered. An example of the dependence of the distribution of Tajima’s *D* on the frequency of the marker is shown in Figure S2.

The results of the previous section show precisely how Tajima’s *D* test values for a fully linked locus is biased as a function of the frequency of the marker. These biases can be computed analytically for all Tajima’s *D*- like SFS-based tests [13] in a similar way, using equation (14) and simple approximations (e.g. replacing *S* by its conditional expected value in their denominator).

Furthermore, it is possible to develop versions of Tajima’s *D* and other frequency-spectrum based neutrality tests that take into account the presence of the neutral marker. The simplest approach follows Rafajlović et al. [42] and consists in replacing the neutral spectrum *θL*/*i* by the conditional spectrum *ξ*_*i*│*m*_, where *m* is the count of the marker in the sample. The usual covariance of the spectrum from Fu [4], which appears in the normalisation of the tests, can be replaced by the one derived by Klassmann and Ferretti [43] for the conditional spectrum. We denote the neutral marker by *ϕ* and the allele count of its derived allele by *m*. We use the observed spectrum 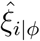 for a window of size *L* containing the marker to build a test of the general form [13, 14]:

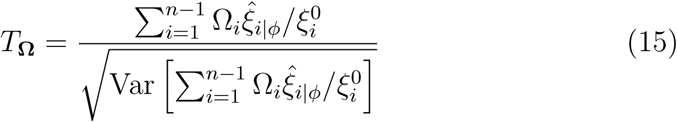

where both the null spectrum 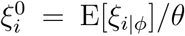 and the variance in the de-nominator are computed under the standard neutral model, i.e. Kingman’s coalescent. The real vector of parameters **Ω** can be chosen in any possible way, as long as it satisfies 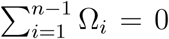. For example, in the absence of a neutral marker, Tajima’s *D* corresponds to 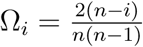.

The definition of the test requires that the null spectrum and the variance are conditioned on the presence of the marker *ϕ*. For the neutral spectrum, it is simply the sum of the expected nested and disjoint spectra presented in equations 14:

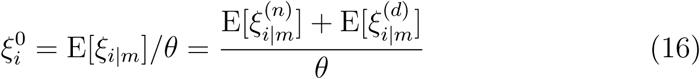

On the other hand, the variance can be decomposed as

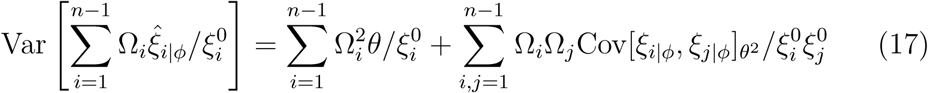

where the *θ* contribution corresponds to the Poisson noise of the mutational process, while the *θ*^2^ contribution to the covariance Cov[*ξ*_*i*│*ϕ*_, *ξ*_*j*│*ϕ*_]_*θ*_^2^ can be easily obtained from the third moments of the spectrum derived by Klassmann and Ferretti [43]:

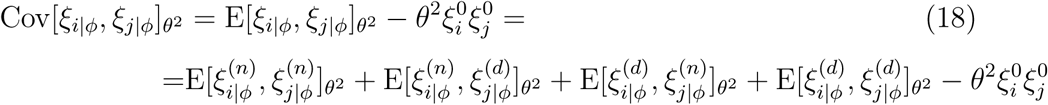

The test can then be built by putting together the results from the last section and an estimation of *θ* and *θ*^2^. The Maximum Composite Likelihood estimate can be used:

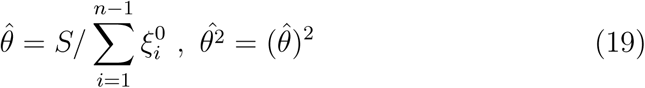

or the Method-of-Moments estimates as in the classical Tajima’s *D*:

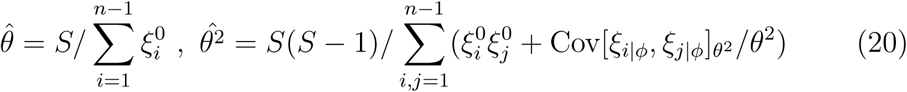

The choice of weights for new tests of this form is someway arbitrary. For example, a modified version of Tajima’s *D* could use the old weights, i.e. 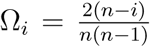, or the old linear coefficients, i.e. 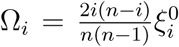, depending if the test should focus on the relative or absolute differences between the null and observed spectrum. Principles and formulae for a meaningful choice of the new weights are discussed in detail by Ferretti et al. [14]. On the other hand, once the weights are chosen, the normalisation does not suffer from any degree of arbitrariness and its form depends only on the third moments computed here.

This straightforward modification of neutrality tests is a promising direction for future dedicated neutrality tests that aim at correcting multiple artefacts such as demography, knowledge of the frequency of the marker, etc.

## 3. Methods

### 3.1 The sample joint 2-SFS

To obtain the sample spectrum for pairs of mutations, we notice that this spectrum can be defined in terms of the expected value of crossproducts of the usual SFS. In detail, we have

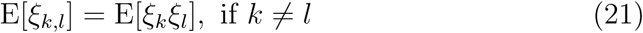

and

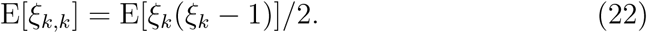

These expected values have been derived by Fu [4] by coalescent methods. However his results do not distinguish the different contributions from nested and disjoint mutations to the spectrum. Tracking the origin of each term in the derivation, it is easy to show that equations (24) and (28) of [4] contribute to nested pairs of mutations, while equations (25), (29) and (30) contribute to disjoint pairs of mutations. All these terms combine linearly and do not interfere, therefore we can decompose the resulting E[*ξ*_*k*_*ξ*_*l*_] into contributions coming from equations (24),(28) and (25),(29) and (30) of [4]. This can be obtained directly by Fu’s expression for the covariance matrix *σ*_*kl*_, since E[*ξ*_*k*_*ξ*_*l*_] = *δ*_*k,l*_E[*ξ*_*k*_] + E[*ξ*_*k*_]E[*ξ*_*l*_] + *θ*^2^*L*^2^*σ*_*kl*_ and E[*ξ*_*k*_] = *θL*/*k*. A detailed review of the calculations of [4], tracking the parts that lead to our mutation classes, is provided in section S3 of the Supplementary Material.

The same results could also be obtained from Theorem 5.1 in Jenkins and Song [34]. In fact, for a special choice of allele transition matrices (in the tri-allelic case, a strictly lower triangular matrix with all non-zero entries equal to 1), their results for recurrent mutations for small *θL* (*θ* in their article) are mathematically equivalent to the results for mutations in an infinite-sites model. Their classification is based on the location of the mutations on the tree: their “nested mutations” correspond to strictly nested and enclosing mutations here, “mutations on the same branch” correspond to co-occurring mutations, “mutations on basal branches” correspond to complementary mutations, and “non-nested mutations’ correspond to strictly disjoint mutations.

### 3.2 The sample conditional 1-SFS

The spectrum for sites linked to a focal mutation of count *l* (equation 13) can be obtained from the previous spectrum (11). The first step is simply to condition on the frequency *l*/*n* of the focal mutation, i.e. dividing the 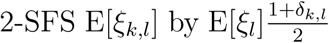 following equations (9) and (10). In fact, 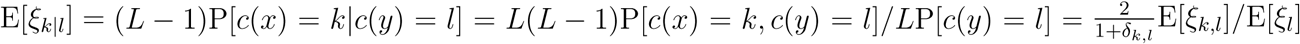 where *c*(*x*) is the derived allele count at site *x*.

The second step is to break further the two contributions of the resulting conditional spectrum into the different components. Strictly nested, cooccurring and enclosing mutations are derived from the nested contribution and are distinguished by site frequencies only: strictly nested ones correspond to *k* < *l*, co-occurring ones to *k* = *l* and enclosing ones to *k* > *l*. Similarly, from the disjoint contribution, mutations belonging to the strictly disjoint component can be obtained by selecting the frequency range *k* + *l* < *n* while complementary ones correspond to *k* + *l* = *n*.

### 3.3 Population spectra

In the limit of large samples, the frequency spectra converge to the continuous SFS for infinite populations. However, the limit *n* → *∞* should be taken with care. The easiest derivation proceeds as follows: since the conditional 1-SFS (eq 14) is a single-locus spectrum, its population components can be obtained from the corresponding ones for finite samples (eq. 13) by direct application of the equation (2). Then the population 2-SFS (eq 12) can be reconstructed from equations (7) and (8), by multiplying by the neutral spectrum E[*ξ*(*f*_0_)] = *θL*/*f*_*0*_ and by 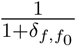 and combining the result into nested and disjoint contributions. The derivation makes use of the following functional limit of the Kronecker delta as a Dirac delta function: *nδ*_⌊*nf*⌋,⌊*nf*0⌋_ → *δ*(*f* – *f*_0_) for *n* → *∞*. More details are given in section S4.

## 4. Discussion

In this article, we have provided exact closed formulae for the joint 2-SFS as well as the first expressions for the conditional 1-SFS, both for sample and population. The 2-SFS was already derived in different forms [34, 37, 36], but the expression presented here for the infinite-sites model is embedded in the framework of Fu [4]. Sample spectra were then used to derive the population spectra by letting *n* → *∞*. Importantly, our results only hold when there is no recombination, and are averaged across the tree space.

The analytical expressions provided in this paper can be intuitively understood in terms of the evolution of linked mutations. Consider a new mutation increasing in frequency by neutral drift and reaching low/intermediate frequency. We expect to find a large number of strictly disjoint and a low number of strictly nested linked mutations, since at the time of appearance of the focal mutation most existing mutations were “strictly disjoint”. The spectrum of strictly nested mutations is more skewed towards rare alleles than predicted by the neutral spectrum 1/*f*, since strictly nested mutations evolve inside an expanding subpopulation. On the other hand, the spectrum of strictly disjoint mutations resembles the neutral one but with a slight bias against rare mutations, since they evolved in a slightly contracting subpopulation.

Note that for sequences linked to a mutation close to fixation, co-occurring and complementary mutations dominate. The contrast between the haplotypes produces a strong “haplotype structure”.

Interestingly, conditioning on the presence of a mutation of frequency *f* impacts the length and balance of the coalescent, as apparent from Figure 4. This can be understood as follows. Rare mutations are common in any realisation of the coalescent tree but especially common in the lower branches, therefore they just increase slightly the tree length and the length of the lower branches compared to the unconditioned case. Instead, mutations of intermediate frequency appear mostly in the upper branches of the tree, therefore the presence of such mutations implies higher, more balanced trees. The effect is even stronger for high frequency mutations, which reside only in the uppermost branches, implying highly unbalanced trees.

There are several potential applications of these results. Here we discuss approaches to correct SFS-based neutrality tests taking into account the presence of a neutral marker strongly linked to the genomic region. These corrections are useful in cases when the region has been selected on the basis of evidence from genome-wide association studies or studies of differentiation based on SNP arrays or other (putatively) neutral markers. This is just an example of possible extensions of neutrality tests based on these results. Other applications include the improvement of population genetic inference techniques based on the SFS, such as composite likelihood [*e.g*. 7, 19, 20, 21] and Poisson Random Field methods [16]. These methods use analytical expressions for the SFS for a single site together with approximations of independence between different sites. For sequences with low recombination, methods could be made more rigorous by assuming independence between different *pairs* of sites, while taking pairwise dependence between sites into account through the two-locus SFS developed here.

The spectrum could also be useful for new neutrality tests based on linkage between mutations. Our results lead to a better understanding of the linkage disequilibrium (LD) structure among neutral loci, therefore they can be immediately applied to LD-related statistics, for example to compute average LD across non-recombining neutral loci. As an example, it can be checked numerically that the expected value of *D* between fully linked derived mutations is 0 according to our equations, as expected from LD theory. Furthermore, they can be used to build neutrality tests optimised to detect positive or balancing selection through its effect on the frequency spectrum of linked sites. The spectra presented here could also provide a neutral model for other scenarios, including structural variants or introgressions from different species or populations. Introgressed sequences from close species can be detected as divergent haplotypes in the locus considered, and if introgressions are rare, then the genetic variability within these haplotypes is described by the nested spectrum linked to the introgressed haplotypes.

The SFS presented here is the simplest two-locus spectrum for neutral, non-recombining mutations in a population of constant size. These results could be extended to variable population size using the approach of [9, 34] and to mutations in rapidly adapting populations using the Λ-coalescent approximation and the results of [44]. However, the most interesting extensions would be to consider (a) non-neutral mutations and (b) recombination.

Adding selection to the two-locus SFS would significantly enhance its potential for most of the applications discussed above. The SFS for pairs of selected mutations has been obtained by [32] as a polynomial expansion, but the numerical computation of this expansion is still cumbersome. Given the simplicity of the expression for the single-locus 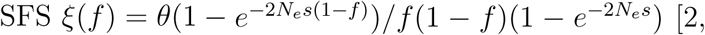 [2, 16], we expect that closed expressions could be found for pairs of mutations with different selective coefficients. This would be a promising development for future investigations.

The classical correspondence between the Kingman model in the large *n* limit and the diffusion approximation suggests that the 2-SFS spectrum presented here is a solution of the diffusion equations for three alleles [3, section 5.10]. In fact, the nested component of the 2-SFS for *f* ≠ *f*_0_ is a stationary solution of the diffusion equation of three alleles of frequency *f*, *f*_0_ – *f* and 1 – *f*_0_:

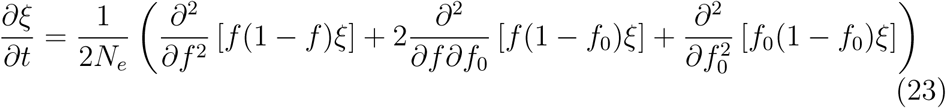

while the disjoint component for *f* ≠ 1 – *f*_0_ is a stationary solution of the diffusion equation of three alleles of frequency *f*, *f*_0_ and 1 – *f*_0_ – *f*:

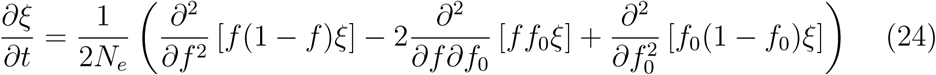

The correspondence implies that the solution (12) is actually the stationary solution of the full set of diffusion equations for the system, including boundary equations for *f* = *f*_0_ and 1 – *f*_0_ and boundary conditions. A direct proof of this result using methods from the theory of partial differential equations could lead to interesting developments towards new solutions for selective equations as well. Our results could also be used to test the accuracy of existing tools based on a numerical solution of the diffusion equations [45].

On the other hand, finding the exact two-locus SFS with recombination appears to be a difficult problem. Recombination is intrinsically related to the two-locus SFS via the same definition of linkage disequilibrium. Obtaining the full two-locus spectrum with selection and recombination could open new avenues for model inference and analysis of genomic data. For this reason, many approximations and partial results have been developed since [25], like expansions in the limit of strong recombination [46]. The SFS of linked loci presented in this paper could be useful as a starting point for different approaches to the effect of recombination events, for example for perturbation expansions at low recombination rates.

An immediate application of our results to recombination events is the following: since in the Ancestral Recombination Graph [47] the recombination events follow a Poisson process similar to mutation events, although with a different rate, the spectrum *ξ*_*k*│*l*_ could also be reinterpreted (up to a constant) as the probability that a single *recombination* event affects *k* extant lineages in a sequence linked to a specific mutation of frequency *l*, i.e. it is equivalent to the spectrum of mutation-recombination events. This approach could be applied to higher moments of the frequency spectrum and lead to new results in recombination theory.

We offer tools for computing the analytical spectra as well as performing simulations by manipulating output of the program *ms* [48]. The corresponding C++ code is contained in the package *coatli* developed by one of the authors and available on http://sourceforge.net/projects/coatli/.

## Acknowledgments

We thank Wolfgang Stephan for insightful discussions. GA and LF were supported by grant ANR-12-JSV7-0007 TempoMut from Agence Nationale de la Recherche. GA was also supported by grant ANR-12-BSV7-0012 Demochips, AK and TW by grants of the German Science Foundation (DFG-SFB680 and DFG-SPP 1590).

## Appendix A. The folded spectra

When no reliable outgroup sequence is available, one cannot assess if the allele is derived or ancestral. In that case, alleles can only be classified as minor (less frequent) and major (most frequent). The distribution of minor allele frequencies, known as the folded SFS, will be noted *η*(*f \**), where *f \** denotes the minor allele frequency that ranges from 0 to 0.5. Importantly, the folded SFS can be retrieved from the full SFS by simply summing alleles at complementary frequencies:

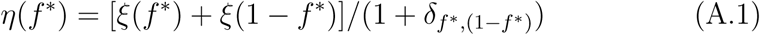

As a consequence, the single site SFS under the standard neutral model then become E[*η*(*f \**)] = *θ*/[*f \**(1 – *f \**)(1 + *δ*_*f*_*\*,*_(1 – *f \**)_)] and E[*η*_*k*_*\**] = *θn*/[*k\**(*n* – *k\**)(1 + *δ*_*k*_*\*,*_*n* – *k*_*\**)], where *k\** denotes the count of the minor allele.

Following the same idea, we define a conditional folded 1-SFS and a joint folded 2-SFS using the minor allele frequencies. Minor alleles can also be classified as “nested” or “disjoint” depending on the presence or absence of individuals enclosing both minor alleles. As for the unfolded case, this classification gives a complete description of the linkage between pairs of mutations. However, in contrast to the unfolded case, the classification has no strict evolutionary meaning. For example, “disjoint” minor alleles do not necessarily correspond to pairs of alleles born in different backgrounds. Moreover, alleles of frequency *f \** = 0.5 (or allele count *k\** = *n*/2) suffer from an ambiguity in the choice of the minor allele and therefore should be treated separately. Note also that with the exception of alleles with frequency 0.5, folded spectra do not contain complementary alleles, since the frequency of one of the two complementary alleles will exceed 0.5.

Pairs of mutations with *f, f*_0_ both larger or smaller than 0.5 will be classified identically (as nested or disjoint) in the folded case. However, pairs of mutations with *f* < 0.5 and *f*_0_ > 0.5 (or vice-versa) will swap their classification. As a consequence, the two components of the 2-SFS are:

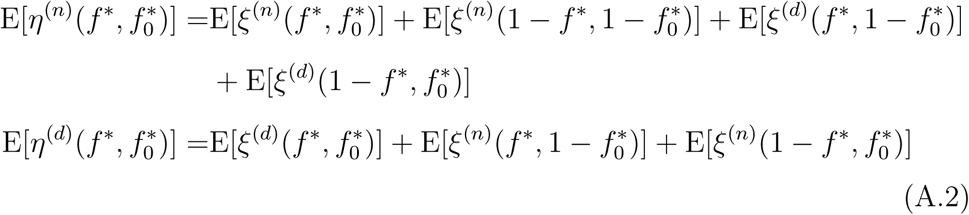

To obtain the conditional 1-SFS, we proceed similarly to the unfolded case. First we separate the 2-SFS above into components based on frequency. The strictly nested component corresponds to frequencies 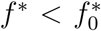 of the nested part, while the co-occurring and enclosing components corresponds to 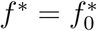 and 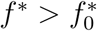 respectively. The strictly disjoint component corresponds to the disjoint part, since there cannot be any complementary component. Then we divide each component by the expected 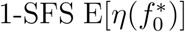 to obtain

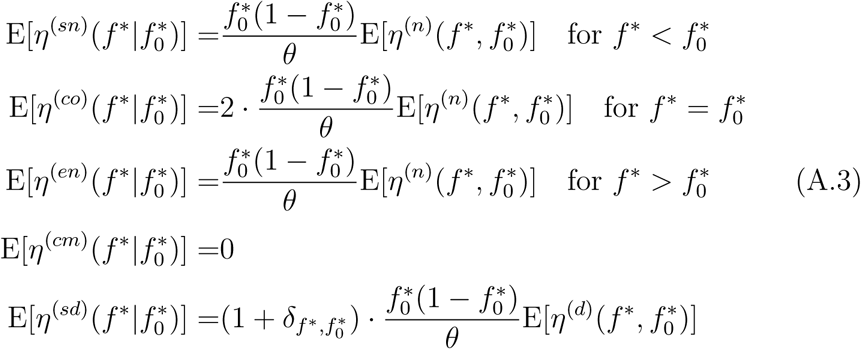

While the classification of the pairs with frequencies *f \** = 0.5 and/or 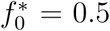 is ambiguous, these pairs are usually irrelevant for the population spectrum.

The sample spectra are similar. For *n* even, there are ambiguous pairs with *k* or *l* = *n*/2 that can be easily retrieved from the equations (11), (13) and treated separately. Considering only *k, l* < *n*/2, the sample 2-SFS is:

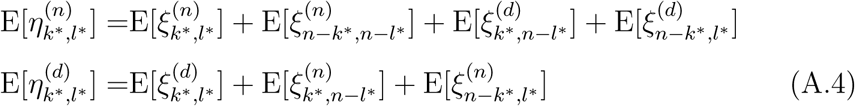

and the conditional 1-SFS is:

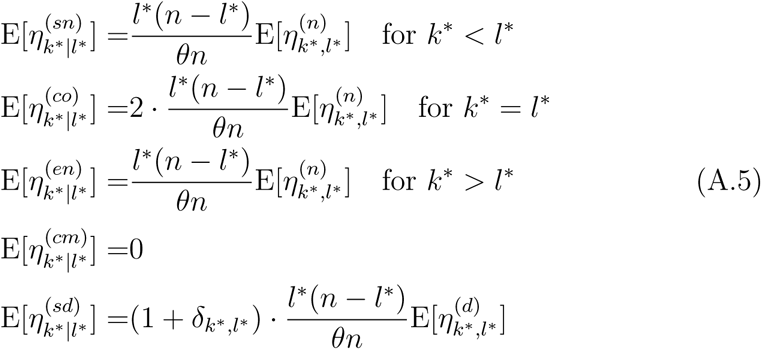

## Footnotes

1 More formally, eq.(2) can be obtained from eq.(1) under the assumptions that 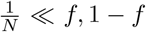, 1 – *f* and that the population SFS is smooth over a range of frequencies 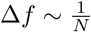.

2 We subdivide the “strictly nested” mutations of [36] into *strictly nested* and *enclosing* mutations while we refer to “identical” mutations as *co-occurring*.

3 Note that the related formula (14) in the paper by [37] has a sign error. It should be identical to the second equation in (11) up to a multiplicative factor.

